# A gene filter for comparative analysis of single-cell RNA-sequencing trajectory datasets

**DOI:** 10.1101/637488

**Authors:** Yutong Wang, Tasha Thong, Venkatesh Saligrama, Justin Colacino, Laura Balzano, Clayton Scott

**Affiliations:** Department of Electrical Engineering and Computer Science, University of Michigan, Ann Arbor, MI 48109 USA; Department of Environmental Health Sciences, University of Michigan School of Public Health, Ann Arbor, MI 48109 USA; Department of Nutritional Sciences, University of Michigan School of Public Health, Ann Arbor, MI 48109 USA; Center for Computational Medicine and Bioinformatics, University of Michigan, Ann Arbor, MI 48109 USA; Department of Electrical and Computer Engineering, Boston University, Boston, MA 02215 USA

## Abstract

Unsupervised feature selection, or gene filtering, is a common preprocessing step to reduce the dimensionality of single-cell RNA sequencing (scRNAseq) data sets. Existing gene filters operate on scRNAseq datasets in isolation from other datasets. When jointly analyzing multiple datasets, however, there is a need for gene filters that are tailored to comparative analysis. In this work, we present a method for ranking the relevance of genes for comparing trajectory datasets. Our method is unsupervised, *i.e.*, the cell metadata are not assumed to be known. Using the top-ranking genes significantly improves performance compared to methods not tailored to comparative analysis. We demonstrate the effectiveness of our algorithm on previously published datasets from studies on preimplantation embryo development, neurogenesis and cardiogenesis.

## 1 Introduction

Recent years have seen rapid growth in the number of single-cell RNA sequencing datasets, giving rise to methods for comparative analyses of these datasets. Examples of such comparative analysis problems include alignment of cells across datasets ^1,2^, assignment of cells in a query to a reference dataset ^3^, and comparison of pseudo-temporal orderings ^4,5^. Due to the high dimensionality of single-cell RNA-seq data, unsupervised feature selection, or gene filtering, is often the first step in many analysis pipelines. However, there are no existing gene filters tailored specifically to comparative analysis. To address this limitation, we develop a gene filter based on the utility of genes for downstream comparative analysis of multiple datasets.

We focus on the comparative analysis task of *data integration*, in the context of datasets where the cells exhibit a trajectory structure. Data integration seeks a common representation of two datasets with the goal of understanding the relationships between the cells across experimental conditions. In particular, the goals of data integration are to remove or reduce batch-effects ^2,1^, i.e., unwanted technical variation between batches, and to maximize mapping accuracy ^3^, a measure of how well cells of the same type are grouped in the shared representation. Our goal is to find a set of features such that when restricted to those features, the two datasets exist on a common trajectory such that cells of the same type are grouped together. When the two batches are obtained from matched biological tissues but sequenced using different techniques, this goal is accomplished by removing batch effects orthogonal to the shared trajectory. Our method also applies more broadly to other settings, including the comparison of datasets obtained from different species and developmental stages.

We introduce *curve-fit objective ranking of gene importance* (CORGI), a novel gene filter that takes as input two datasets and outputs a ranked list of genes based on their potential for integrating trajectory datasets. CORGI is scale-invariant, meaning it produces the same result regardless of the gene expression unit, *e.g.*, raw counts, log counts, or transcripts per million, and the two input batches can be in different units. In addition, CORGI is unsupervised, meaning it requires no knowledge of cell meta-data, while allowing the user to incorporate genes known a priori to be relevant, *i.e.*, marker genes. We demonstrate CORGI on a variety of publicly available datasets and sources of between-batch variation: preimplantation embryo development between mouse ^6,7^ and human ^8^, neurogenesis collected with different cell isolation methods ^9,10^ and cardiogenesis samples collected at different embryonic age ^11,12^.

## 2 Results

The CORGI algorithm ranks genes based on how well they capture the common trajectory in two datasets. The intuition is that a given set of genes *G* is informative of the common trajectory when the pooled data, restricted to only genes in *G*, can be fit by a principal curve with low error. Motivated by this intuition, we repeatedly sample random gene sets, fit a principal curve to the filtered data, and keep a running average of the fit error for all the genes (Figure 1B). After running the random sampling for a user-specified amount of time, we rank the genes based on the average of the fit error and keep genes with low error in the gene filter (Figure 1C). For the mathematical details, see Section 6.5.

**Figure 1:**
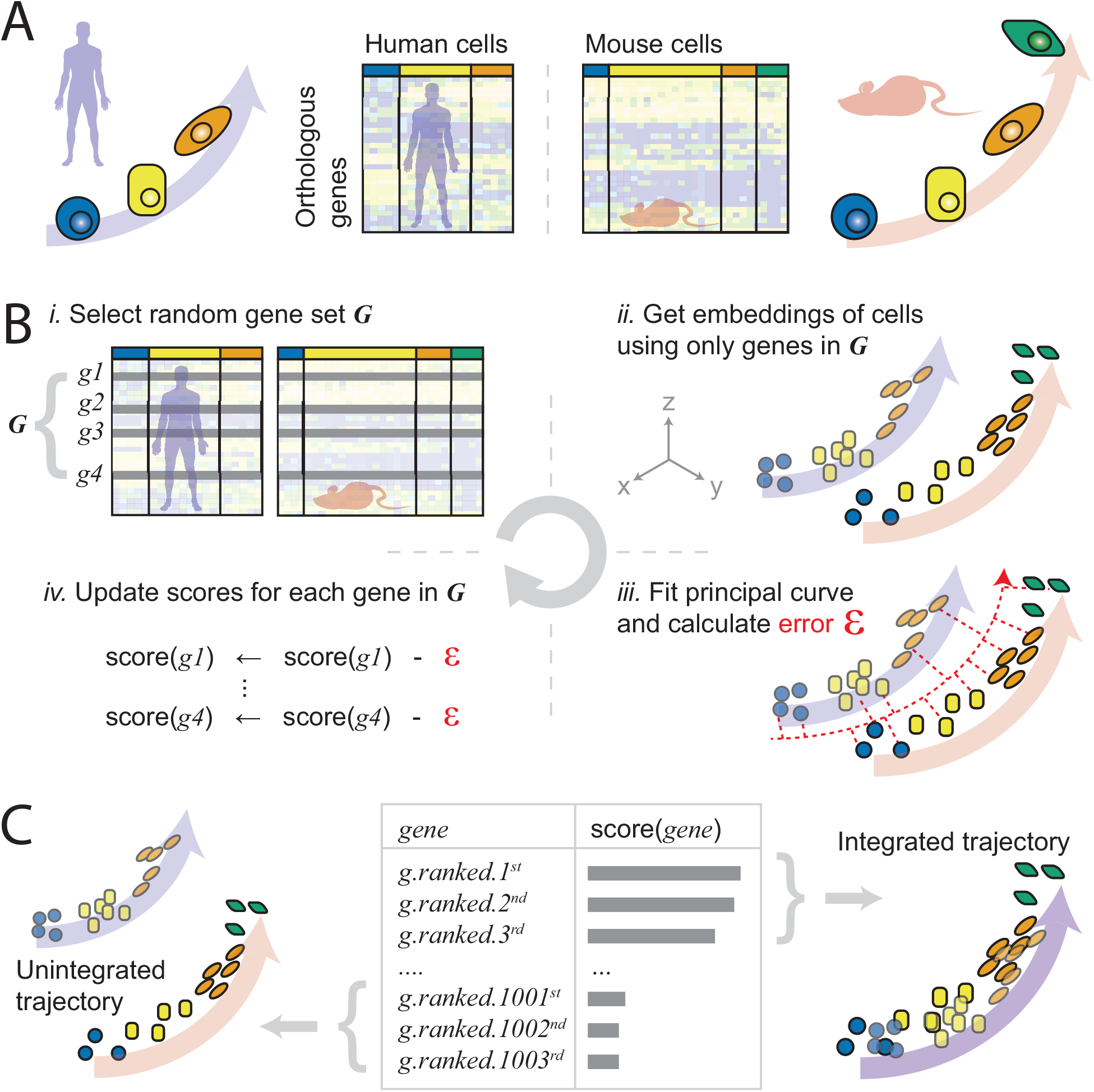
Overview of the CORGI algorithm. A. CORGI takes as input a pair of single-cell gene expression matrices. B. The main loop of the CORGI algorithm. The trajectory integration score of each gene is initialized to zero. B.i the main loop of CORGI starts by selecting a random gene set. B.ii a low dimensional representation of the cells of the combined dataset is computed. B.iii a principal curve is fit to the low dimensional representation and the mean square error of the fit *ϵ* is computed. B.iv the trajectory integration score for every gene in the random gene set is incremented by *ϵ*. C. CORGI outputs a ranked list of genes where the set of the highest ranked genes integrates the two trajectories.

Although CORGI takes a randomized approach, we observe that multiple independent runs have similar output (Figure S7 and S6). Furthermore, we find that the embeddings of the cells produced using CORGI gene sets is robust to the number of top ranking genes selected (Figure 2 and S4, using top 100 and 500 genes, respectively).

**Figure 2:**
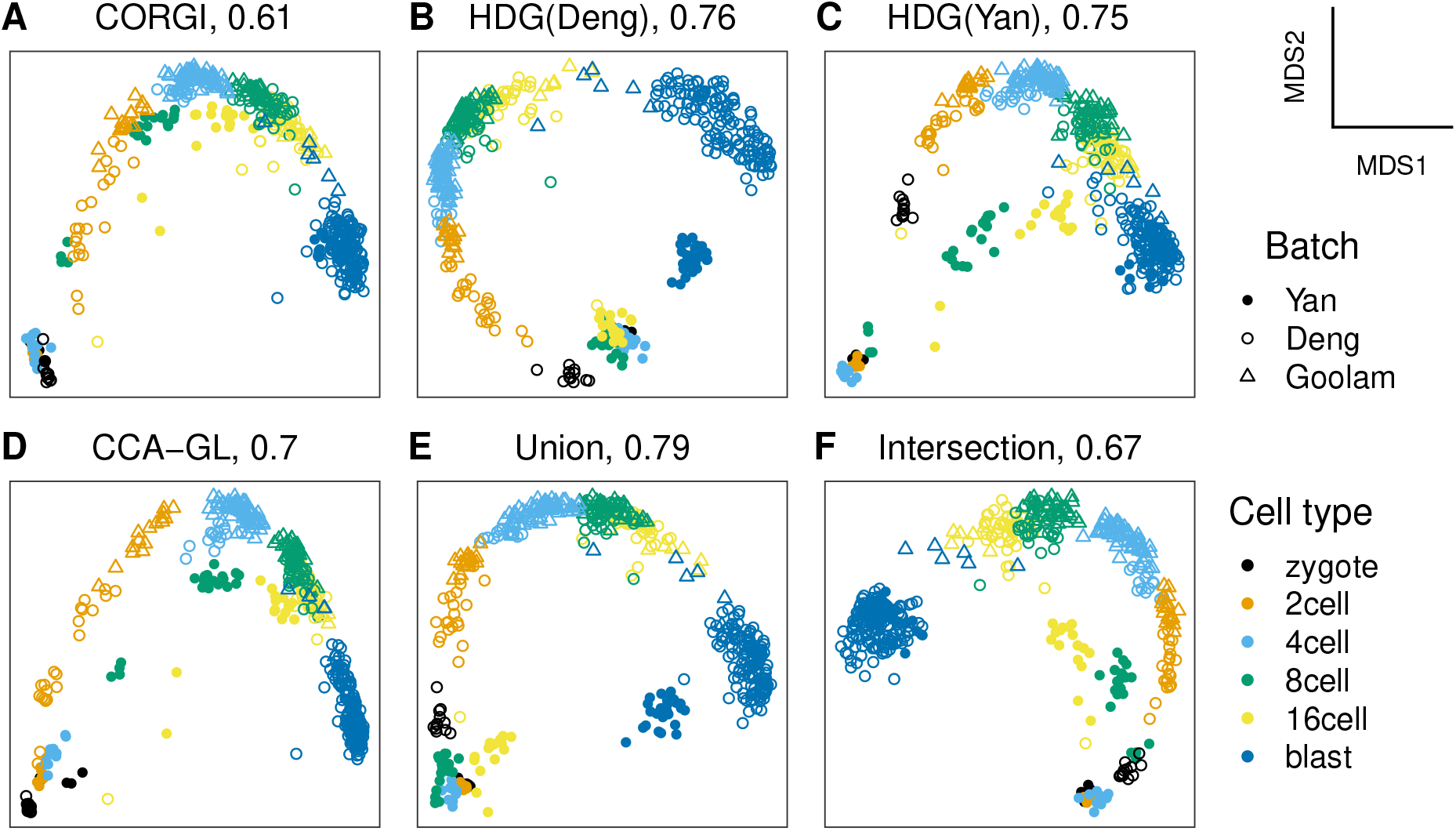
Preimplantation embryogenesis in human ^8^ and mouse ^6^. A-F, The title of each scatterplot denotes the shortened name of the gene set and the batch separation metric. B and C, highly dropped-out genes of the two respective batches. D, genes with the highest canonical correlation analysis gene loading scores. E and F, union and intersection of highly dropped-out genes of the two datasets, respectively.

For evaluation criteria, we consider batch separation score, mapping accuracy and visual quality of two-dimensional embeddings of the integrated datasets. The embedding is obtained from multidimensional scaling ^13^ applied to the distance matrix derived from Spearman *ρ* correlation. See Methods for details and justifications.

### 2.1 Preimplantation embryo development between mouse and human

Preimplantation embryogenesis follows a continuous trajectory but forms discrete clusters — zygotes, 2-cells, 4-cells and so on — along the trajectory. However, comparing embryogenesis between human and mouse isn’t completely straightforward. For instance, the clusters, or the developmental stages, between species do not necessarily match up, *e.g.*, the gene expression pattern of a human 16-cell embryo may more closely resemble that of a mouse 8-cell embryo than a 16-cell embryo. Furthermore, certain transcriptional programs may be species-specific ^14^. Therefore, it is of interest to find a common set of genes that capture the transcriptional dynamics during embryogenesis.

We apply CORGI to human and mouse preimplantation embryo development datasets from Yan et al. ^8^ and Deng et al. ^6^, respectively. We compare against another five gene filters. The first four are highly dropped-out genes: *HDG(Mouse)*, *HDG(Human)*, their *union*, and their *intersection*. The fifth gene filter, *CCA-GL*, consist of genes with the highest canonical correlation analysis gene loading scores ^1^. See Section 6.2 for details. Below, gene set names are italicized, *e.g.*, *CORGI* or *Intersection*. All gene sets have 100 genes. See Supplementary Information Section 4.1 for the genes.

In Figure 2, we plot the results of MDS on the gene sets considered in this section. The format of the plot title, *e.g.*, “CORGI, 0.53”, is the name of the gene set used followed by the *batch separation metric*, where larger number implies stronger batch effects. See Section 6.3.1 for detail on the definition of the metric. Although we select genes using only the datasets from Deng et al. ^6^ and Yan et al. ^8^, we visualize additionally a third mouse preimplantation embryo dataset from Goolam et al. ^7^ In Figure 2, the mouse and human cells are represented by open and solid shape markers, respectively. We note that *CORGI* achieved the lowest batch separation metric, which agrees with the visual observation that only *CORGI* is able to integrate the mouse and human datasets into a single trajectory.

### 2.2 Neurogenesis datasets with different cell isolation methods

Often there is a lack of consensus on the method for isolating a population of cells of interest from their niche, *e.g.*, isolation of adult neural stem cells ^15,16^. Datasets generated by different labs might use different isolation methods and therefore may suffer from batch effects. Thus, it is of interest to find gene sets reflecting interesting biology common to the two populations, rather than technical differences.

We apply CORGI to two datasets on neurogenesis in the subventricular zone (SVZ) produced by Llorens-Bobadilla et al. ^9^ and Dulken et al. ^10^ The two datasets were obtained useing different surface proteins in fluorescence-activated cell sorting to purify neural stem cell (NSC) and neural progenitor cell (NPC) populations. Furthermore, Dulken et al. ^10^ used supervised feature selection to rank genes that best distinguish between the NSC and NPC populations. Their method output a gene list *Cons_n_* of top *n* ranked genes, where they selected *n* = 34. Here, we run CORGI with *Cons*_34_ as marker genes. For the final gene set, we merge the top 100 CORGI genes with *Cons*_34_, resulting in 134 genes in total. See Section 6.5 for detail on using marker genes with CORGI. To maintain fairness, we augment all compared gene sets as discussed in Section 6.2 with *Cons*_34_. Furthermore, we compare a fully supervised gene filter by considering the gene set *Cons*_134_, *i.e.*, the top 134 genes from the ranked list by Dulken et al. ^10^ which also contains *Cons*_34_.

In Figure 3, we use MDS to visualize the two datasets, where, following Dulken et al. ^10^ we exclude the astrocytes as they are indistinguishable from the early NSCs. From Figure 3A-G, only *CORGI*, *CCAGL* and *Intersection* are able to remove the batch effects from the MDS plots. Although, *CCA-GL* and *Intersection* achieve a lower batch separation metric than *CORGI*, visual inspection suggests that *CORGI* more sharply distinguishes between the NSCs and the NPCs. To quantify this, we benchmark the mapping accuracy capability of each of the gene sets using scmapCluster ^3^ in Figure 3G. The output of scmapCluster is benchmarked by Cohen’s *κ* where a larger number implies higher mapping accuracy. For the definition of mapping accuracy, see Section 6.3.2. The *x*-axis of Figure 3G shows the threshold parameter of scmapCluster used.

**Figure 3:**
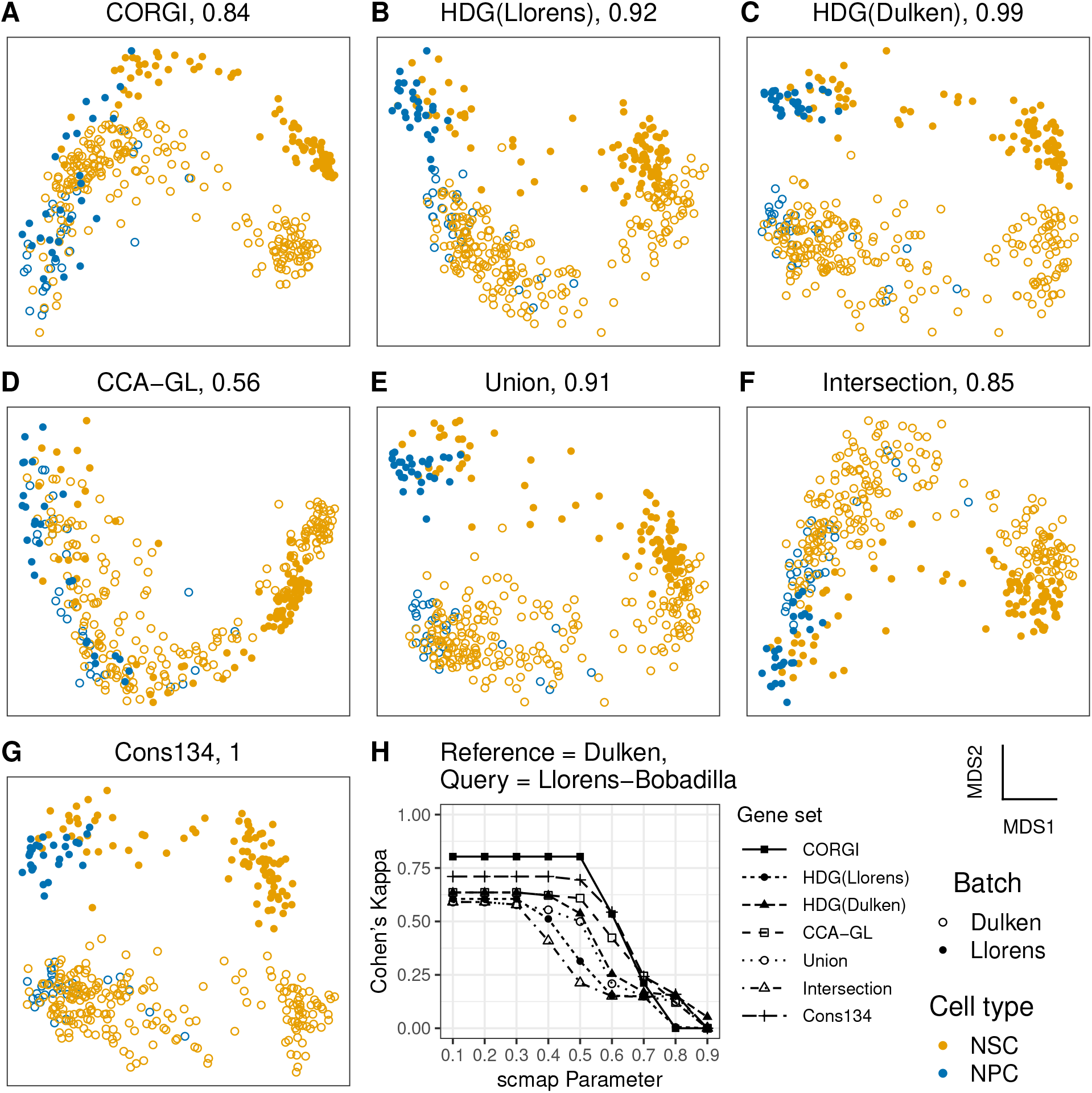
Neurogenesis in the subventricular zone. A-G, The title of each scatterplot denotes the shortened name of the gene set and the batch separation metric. B and C, highly dropped-out genes of the two respective batches. D, genes with the highest canonical correlation analysis gene loading scores. E and F, union and intersection of highly dropped-out genes of the two datasets, respectively. G, top genes according to the supervised method in ^10^. H, scmapCluster prediction of cell-types in query using the reference dataset. X-axis is the threshold hyperparameter used for scmapCluster. Y-axis shows the mapping accuracy between the ground truth and the prediction for cells in the query dataset.

For the thresholds between 0.1 and 0.7, *CORGI* achieved the highest *κ* (Figure 3G). For thresholds greater than and equal to 0.7, *κ* drops below 0.25 for all gene sets, suggesting that the proper choice of threshold should be less than 0.7. Notably, *CORGI* is able to map the cell types more accurately across the two datasets than even the fully supervised *Cons*_134_ gene set. This suggests that genes informative for explaining cell types in one dataset may not generalize well to a new dataset.

### 2.3 Cardiovascular progenitors

Comparing time course experiments where samples are collected at different developmetal stage or ages adds an additional layer of subtlety to the analysis. We consider two published mouse embryonic developmental datasets by Scialdone et al. ^11^ consisting of epiblast cells from embryonic day 6.5 (E6.5) and mesodermal cells from E7.5, and by Lescroart et al. ^12^ consisting of cardiovascular progenitors from E6.75 and E7.25. As shown by Lescroart et al. ^12^, the finer time course scRNAseq data from E6.75 and E7.25 bridge the gap between E6.5 and E7.5 forming a smooth developmental trajectory. We apply CORGI to the Scialdone and Lescroart datasets to find the top 100 genes that best reflect this developmental trajectory. See Supplementary Information Section 4.3 for the genes.

In Figure 4A, the panel corresponding to *CORGI*, the trajectory structure is clearly visible. The other filters that somewhat reveal a trajectory, *e.g.*, *CCA-GL*, do not represent the temporal structure as well as *CORGI*. In contrast to the previous two sections, batch separation is desirable, since the embryonic age of the cells across the two batches do not non-overlap. More specifically, cells from the Scialdone dataset are situated at the endpoints of the trajectories while the Lescroart dataset interpolates the endpoints, forming a bridge. The plot produced using CORGI genes best demonstrates the temporal relationships in this developmental trajectory.

**Figure 4:**
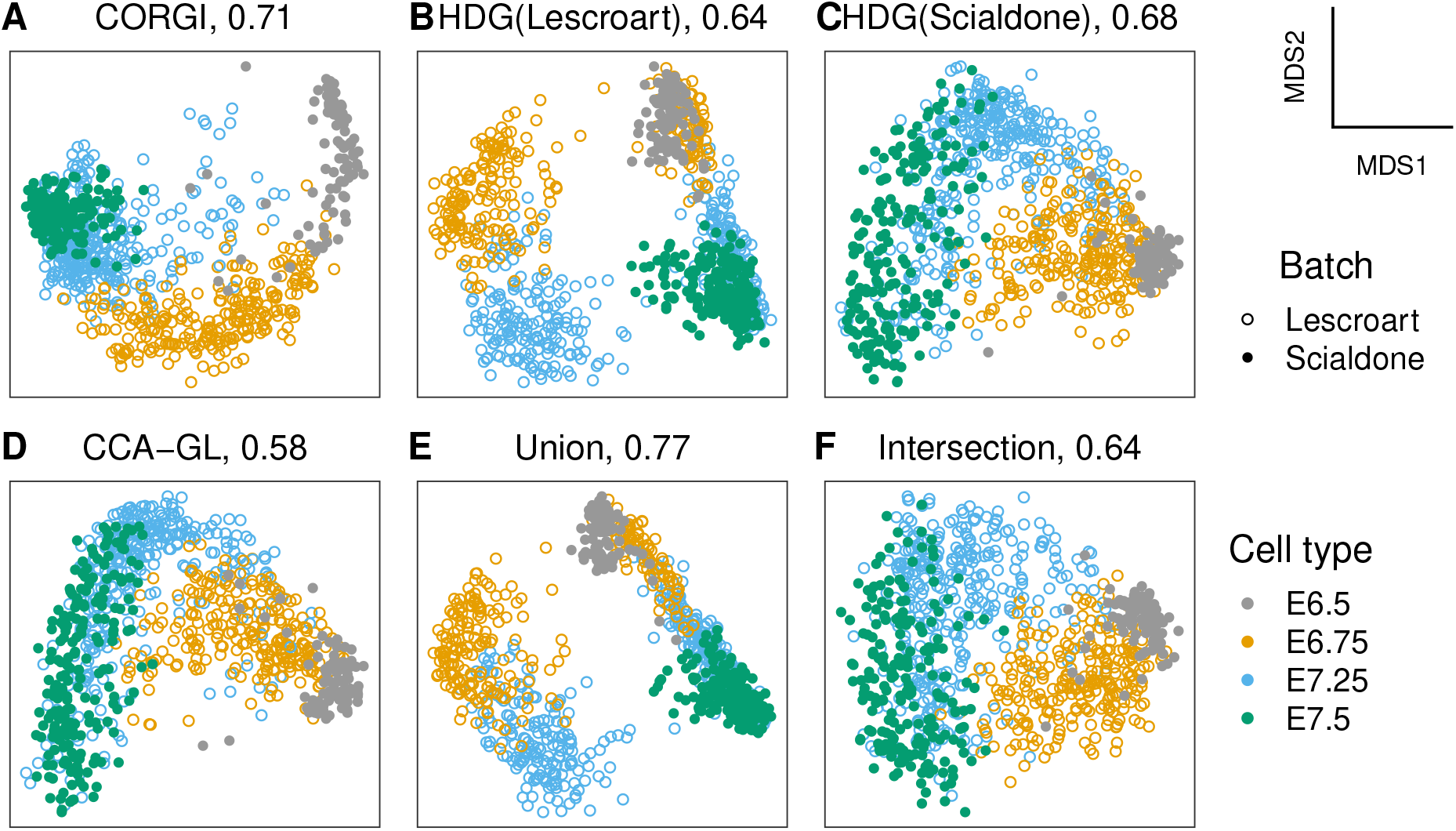
Cardiogenesis. A-F, The title of each scatterplot denotes the shortened name of the gene set and the batch separation metric. B and C, highly dropped-out genes of the two respective batches. D, genes with the highest canonical correlation analysis gene loading scores. E and F, union and intersection of highly dropped-out genes of the two datasets, respectively.

We note that the Lescroart dataset consists of two replicates. Thus, there are batch effects internal to the Lescroart dataset that are visible in the plot generated by highly dropped-out genes (Figure 4B and Figure S8). Even in the union of *HDG(Scialdone)* and *HDG(Lescroart)*, this internal batch effect can be clearly observed, suggesting that the variation explained by the internal batch effect is dominant. In contrast, the internal batch effects are not seen in plots generated with *CORGI, CCA-GL, Intersection, HDG(Scialdone)* (Figure 4A, C, D, and F).

## 3 Discussion

We have shown that the choice of gene filter has a dramatic impact on data integration and that CORGI is useful for integration across differing species, tissue dissociation techniques, and time courses. Our approach is tailored to integrating trajectory dataset and therefore can find genes more relevant to the developmental processes of interest. Improved gene filters for integration may impact biological understanding, allowing us to draw new biological insights from where cells map relative to each other via pseudotime analyses. Applying CORGI on paired experiments with differing conditions can potentially identify genes informative of the shared heterogeneity, *e.g.*, evolutionarily conserved pathways in preimplantation embryogensis. The integrated analyses can be used to map one poorly characterized organism to a better characterized one. We are hopeful that CORGI can be applied in the future to elucidate less well-characterized biological systems. Given the exploratory and high-throughput nature of scRNAseq experiments, CORGI may provide a compact but informative panel of genes useful for lower-throughput experiments such as spatial transcriptomics ^17^.

## 4 Author Contributions

Y.W. and C.S. wrote the paper. Y.W. designed the algorithm and implemented the software with the help of V.S., L.B., and C.S. Y.W. performed the data analysis with the help of T.T. and J.C.

## 5 Acknowledgments

J.C. and T.T. were supported in part by NIH/NIEHS Grants R01 ES028802 and P30 ES017885. L.B. was supported in part by ARO YIP award W911NF1910027, DARPA grant 16-43-D3M-FP-037. L.B., C.S., and Y.W. were supported in part by NSF BIGDATA award IIS-1838179. C.S. and Y.W. were supported in part by the Chan Zuckerberg Initiative (grant number 2018-183157). L.B., J.C., C.S., T.T., and Y.W. were supported in part by the Michigan Institute of Data Science (MIDAS) grant for Health Sciences Challenge Award.

## 6 Methods

Let *X*_1_ and *X*_2_ be the gene expression matrices corresponding to batch 1 and 2 respectively. For *b* ∈ {1, 2}, *X_b_* is the gene-by-cell count matrix with *d* rows (genes) and *n_b_* columns (cells). Below, we use *G* to denote a generic gene set *G* ⊆ {1, 2*, …, d*}.

### 6.1 Dimensionality reduction

For dimensionality reduction, we use multidimensional scaling (MDS) on the distance matrix derived from the Spearman *ρ* correlation, which is invariant to monotone transformation of the data. The details are explained in the next paragraph. Other dimensionality reduction methods considered are principal component analysis ^13^ (Figure S1), canonical correlation analysis ^1^ (Figure S2), and PCA on the mutual nearest neighbor batch-corrected data ^2^ (Figure S3). MDS does not require normalization and visual inspection suggests that MDS performs better.

To compute a cell-cell distance matrix, the columns (cells) of *X*_1_ and *X*_2_ are first pooled together to form *Y* = [*X*_1_*, X*_2_] and row-downsampled to genes in *G*. Denote the resulting |*G*|-by-(*n*_1_ + *n*_2_) matrix by *Y_G_*. We compute the Spearman rho correlation matrix *R* between all pairs of columns of *Y_G_*, *i.e.*, *R* is A square matrix of width *n*_1_ + *n*_2_. Taking 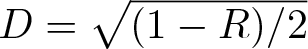 converts the correlations into distances ^18^. Finally, classical multidimensional scaling ^13^ is run on *D*.

### 6.2 Comparison with existing gene filters

The *highly variable gene* ^19^ (HVG) filter selects genes with higher than expected *coefficient of variation*. Kiselev et al. ^3^ recently proposed an analogous method using *dropout rate* rather than coefficient of variation. We refer to the dropout-based method as the *highly dropped-out gene* (HDG) filter. In Supplementary Figure S5, we compared HDG and HVG and found that HDG performed better. Therefore, in this work, we compare only against HDG.

The HDG filter is designed for a single batch of data. We consider two extensions of HDG to the two-batch setting: the *union HDG*(*X*_1_) ∪ *HDG*(*X*_2_) and the *intersection HDG*(*X*_1_) ∩ *HDG*(*X*_2_). To ensure fair comparison, all gene sets are kept the same size. For details, see Supplementary information Section 3.

We also consider a gene filter where genes are ranked according to the gene loading scores from canonical correlation analysis ^1^ run on *X*_1_*, X*_2_ using the default parameters, abbreviated as *CCA-GL*.

When marker genes *M* ⊆ {1*, …, d*} are available, all gene sets are augmented by *M* . See Supplementary Information Section 3.2. All gene sets obtained are listed in Supplementary Information Section 4.2.

### 6.3 Criteria for comparing gene sets

We consider two metrics: *batch separation*, which is always computable, and the *mapping accuracy*, which is computable only when cell-type labels are available for both batches. Batch separation measures how easily the batch labels can be recovered from the scatter plot. The mapping accuracy measures how tightly clustered are the cells of the same type across both batches.

#### 6.3.1 Batch separation

We use the svm function with default parameters in the e1071 R library with the 2D coordinates of the cells as training data *x_train_* and the batch labels as the response variable *y_train_*. The coordinates are normalized to have unit variance in each component. An SVM with default parameters is trained on *x_train_, y_train_*. Using the trained model, we predict the batch labels *y_predict_*. The *batch separation score* is defined as the Cohen’s *κ* between *y_predict_* and *y_train_*. When *κ* is closer to 1, batch label can be accurately predicted, implying stronger batch effects. Conversely, when *κ* is exactly zero, the batch labels can not be predicted better than random guessing.

#### 6.3.2 Mapping accuracy

To quantify mapping accuracy, we reserve one batch with its cell-type labels *ℓ_r_* as the *reference* dataset to predict the cell-type labels *ℓ_q_* of the other batch, the *query* dataset. We use the scmapCluster ^3^ algorithm to match the cells in the query dataset to that of the reference dataset, producing an estimate 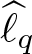 of *ℓ_q_*. Following Kiselev et al. ^3^, we use Cohen’s *κ* as a measure of agreement between the estimate 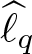 and the truth *f_q_*. The scmapCluster algorithm takes a threshold hyperparameter, for which we sweep over the range {0.1, 0.2*, …,* 0.9}.

### 6.4 Measuring trajectory integration score of a gene set

The core component of CORGI is the *trajectory integration score* that measures the capacity of a gene set for capturing the shared trajectory structure in two datasets. CORGI produces a ranking of all genes by this score. In the previous section, we discussed batch separation and mapping accuracy as benchmarking metrics. However, we argue below that neither of these metrics is an appropriate trajectory integration score.

Batch separation is not an adequate trajectory integration score because selecting genes that have zero expression collapses both datasets to a single point, resulting in low batch separation. However, such a gene set is uninformative. In cases when the developmental stages are distinct, *e.g.*, the cardiogenesis datasets ^11,12^, it is even undesirable to mix the batches together.

Mapping accuracy is also not an adequate trajectory integration score, because first and foremost computing mapping accuracy requires cell-type labels. Even when labels are available, the batches may not have any overlapping cell-types. When the cell-types from the two datasets do not overlap at all, it may still be sensible to perform data integration. This is the case in the datasets from Scialdone et al. ^11^ and Lescroart et al. ^12^, where the cells are collected from different embryonic developmental times. Even when labels are available, it may be unclear how the cell-types match up between the batches, *e.g.*, comparing human and mouse preimplantation embryo development^6,8^. Therefore, we should not design the score to penalize mismatches.

The objective value of a curve-fit algorithm, *e.g.*, the principal curve ^20^, offers a way to measure the trajectory integration capability of a gene set, assuming only that the two batches of data occupy a single continuous trajectory. Another way to state the assumption is that when filtered to a gene set that is relevant to the trajectory, both datasets are explained by a single trajectory.

In contrast to gene filters that rank genes based on statistics derived from a single gene, CORGI ranks the genes by testing several genes in combination. Thus, another factor in CORGI’s success is that it scores genes according to their informativeness when combined with other genes. This is in contrast to filters liike HVG and HDG which consider genes in isolation.

### 6.5 Mathematical details for CORGI

*Input*. *X*_1_ and *X*_2_ data matrices for the two batches.

*Parameters*. Default values are shown in parentheses.

- *r* (200) the size of the random gene set,
- *k* (2) the dimension of the latent embedding of the cells,
- *T* (2 hour) the wall-clock time budget for the algorithm.

*Optional parameter*. A set of markers genes *M* ⊆ {1*, …, d*} can be provided. By default, *M* = ∅.

*Initialization*. For each non-marker gene *g* ∈ {1, 2*, …, d*} \ *M*, let *c_g_* denote the CORGI score which is initialized to zero for all genes. Furthermore, let *N_g_* denote the number of times that gene *g* has been sampled, also initialized to zero.

*Main loop*. We first uniformly sample a random set *G′* ⊆ {1, 2*, …, d*} \ *M* of genes of a fixed size |*G′*| = *r*. Let *G* = *G′* ∪ *M*. We compute the trajectory integration score *s*(*G*) to be defined later. For each *g* ∈ *G*, perform the update *C_g_* ← *C_g_* + *s*(*G*) and *N_g_* ← *N_g_* + 1. While the running time of the algorithm has not exceeded the budget *T*, the loop is repeated.

*Output*. The normalized CORGI scores 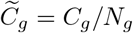 for each *g* ∈ {1*, …, d*} \ *M* where genes with higher 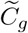 are more informative of the common trajectory in *X*_1_*, X*_2_.

*Trajectory integration score*. Here, we detail the computation of the trajectory integration score *s*(*G*) for a given gene set *G* ⊆ {1, 2*, …, d*}. Consider the *rank correlation matrix*

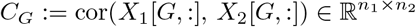

whose *i, j*-th entry is the Spearman correlation coefficient between *X*_1_[*G, i*] and *X*_2_[*G, j*]. When *n*_1_ and *n*_2_ are larger than 200, we further downsample the cells to randomly sampled sets *S_b_* ⊆ {1*, …, n_b_*} where |*S_b_*| = 200. Subsequently, we use the downsampled correlation matrix

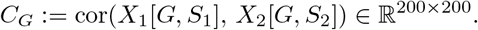

The sets *S*_1_*, S*_2_ are randomly resampled during every iteration of the loop. Define the similarity matrix *W_G_* := (1 + *C_G_*)*/*2 whose entries capture similarity between cells in different batches.

We compute the *Z* matrix as defined by Dhillon ^21^ for the similarity matrix *W_G_*. Similar to canonical correlation analysis, *Z* is an *n*_1_ + *n*_2_ by *k* matrix that embeds of the cells in *k*-dimensional space. To be self-contained, we outline the computational steps of *Z*, while referring the interested reader to the original paper by Dhillon ^21^ for the theoretical motivation of the method.

The computation of *Z* is identical to canonical correlation analysis ^1^. In fact, *Z* is canonical correlation analysis performed on the rank-transformed data. Define *D_r_* = diag(rowSums(*W_G_*)) to be the diagonal matrices whose diagonal entries are the of the row sums of *Z*. Analogously define *D_c_* = diag(colSums(*W_G_*)) with columns instead of rows. Let 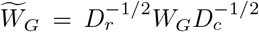. Compute the singular value decomposition 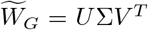 where the columns of *U* = [*u*_1_*, u*_2_*, …*] and *V* = [*v*_1_*, v*_2_*, …*] are ordered by the magnitude of the singular values from largest to smallest. Take *U_k_* = [*u*_2_*, …, u_k_*_+1_] and *V_k_* = [*v*_2_*, …, v_k_*_+1_]. The matrix *Z* is taken to be

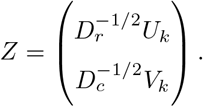

The principal curve algorithm ^20^ is applied to *Z*, fitting a curve to model the assumption that the data come from a continuous trajectory. Finally, we compute *ϵ*, the mean-square error of the curve-fit, and put *s*(*G*) = −*ϵ*.

It is important that the batch effects are removed before curve fitting when computing the embedding *Z* in the CORGI main loop. Otherwise, suppose *Z* perfectly separates the two batches where each batch clusters tightly about its center, then choosing a principal curve that connects the two cluster centers results in low fitting error. However, such a embedding *Z* is not informative for comparative analysis. Hence, we use the method of Dhillon ^21^ for computing *Z* rather than the visualization method described in Section 6.1. The method of Dhillon ^21^ removes batch effects because it performs CCA on rank-transformed data.

### 6.6 Software availability

CORGI is open-source and available as an R package on GitHub (https://github.com/YutongWangUMich/corgi). Code used to generated figures in this paper and supplement is also available (https://github.com/YutongWangUMich/corgifigures).

