## Supplementary Information for "A gene filter for comparative analysis of single-cell RNA-sequencing trajectory datasets"

### 1 Supplementary information

#### 1.1 Public datasets analyzed

| *Dataset* | *Reference* | *Accession code* |
|---|---|---|
| Human preimplantation embryo | Yan et al.[7] | GSE36552 |
| Mouse preimplantation embryo | Deng et al.[1] | GSE45719 |
| Mouse preimplantation embryo | Goolam et al.[3] | E-MTAB-3321 |
| Neurogenesis in the SVZ | Dulken et al.[2] | PRJNA324289 |
| Neurogenesis in the SVZ | Llorens-Bobadilla et al.[5] | GSE67833 |
| Mouse cardiovascular progenitor cells | Lescroart et al.[4] | GSE100471 |
| Mouse mesoderm | Scialdone et al.[6] | E-MTAB-4079<br>E-MTAB-4026 |

#### 2 Supplementary figures

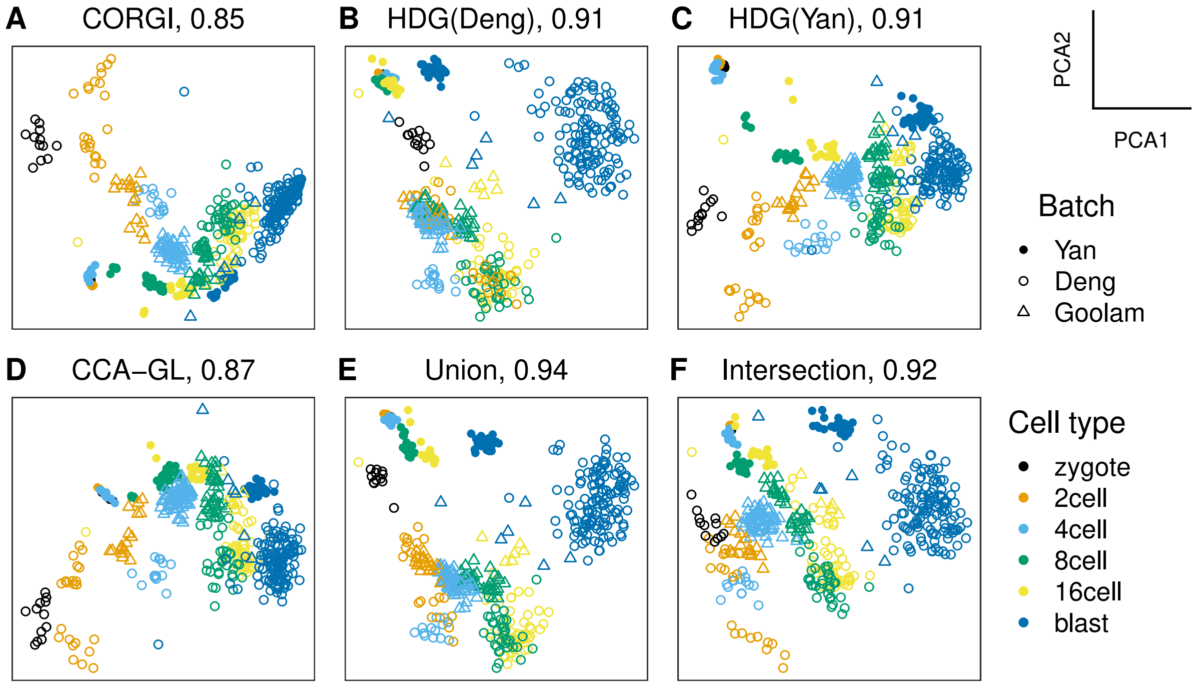


Figure S1: PCA on the embryogenesis datasets across various gene sets. Related to Figure 2 in the paper. The temporal structure of the dataset is not well represented compared to Figure 2.

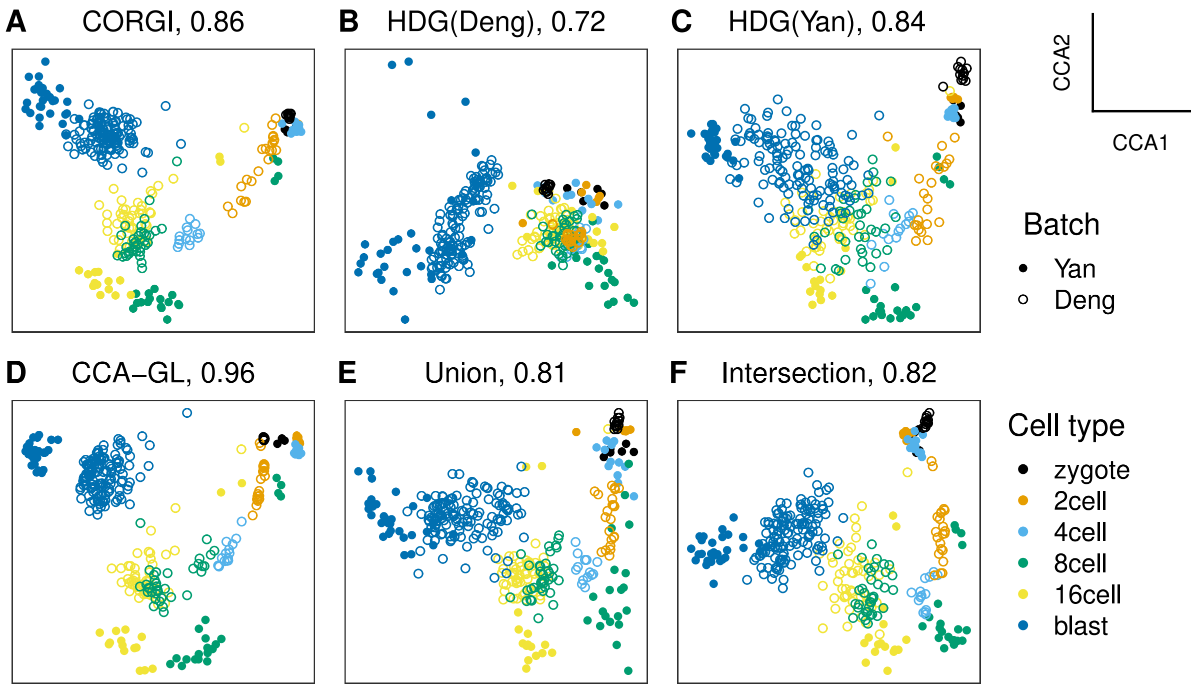


Figure S2: Canonical correlation analysis dimensionality reduction on the embryogenesis datasets across various gene sets. The temporal structure of the dataset is not well represented compared to Figure 2.

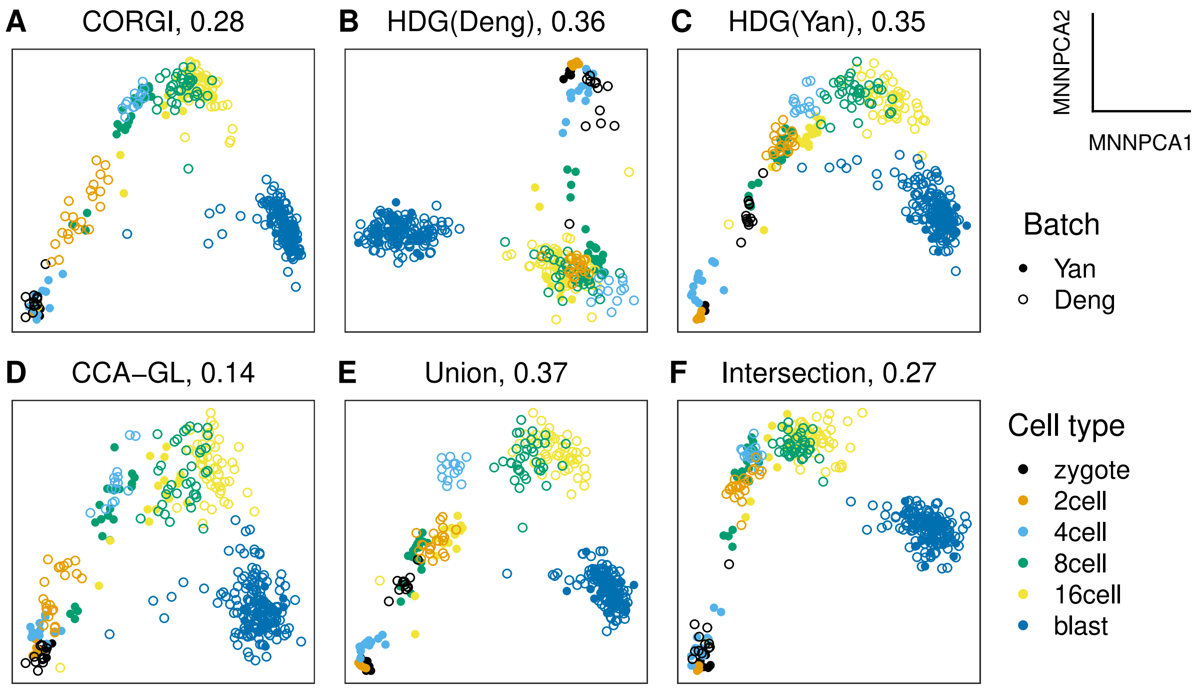


Figure S3: PCA on the mutual nearest neighbor batch-corrected embryogenesis datasets across various gene sets. The temporal structure of the dataset is well represented, similar to Figure 2. Furthermore, the batch separation score is low due to the explicit batch correction employed by the algorithm. However, in our main paper, we show multidimensional scaling since it does not explicitly correct for batch effects. This allows us to examine in isolation the impact of the gene filters on batch effect removal.

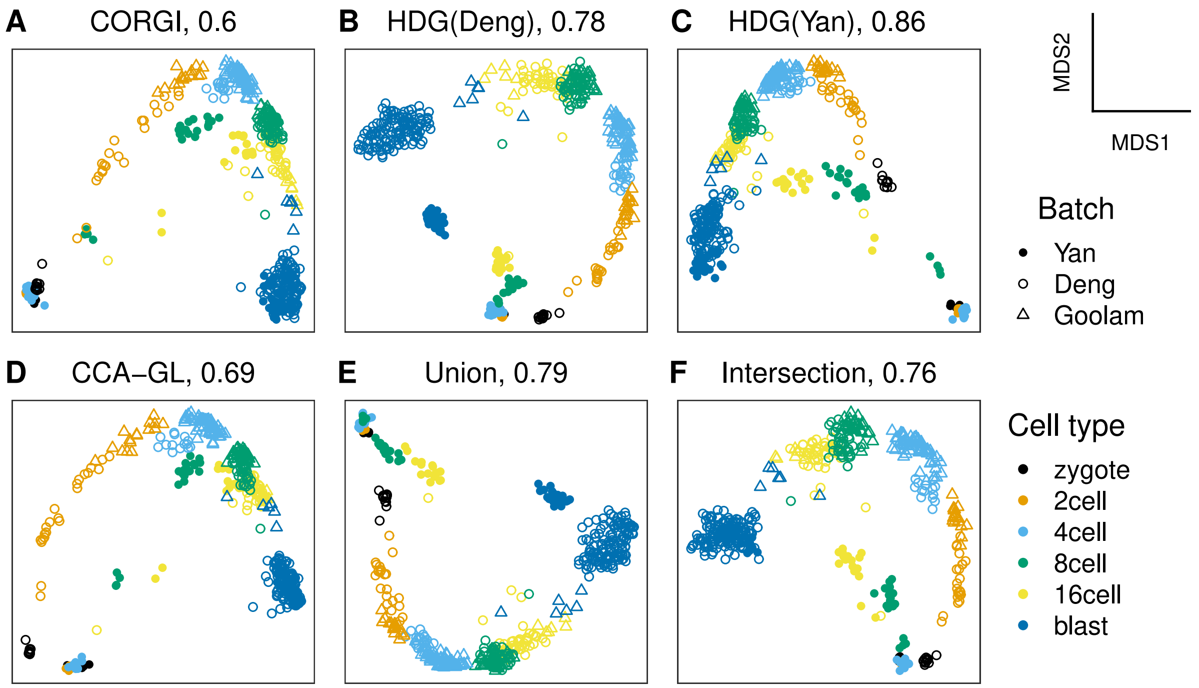


Figure S4: MDS on the embryogenesis datasets across various gene sets. Similar to Figure 2 in the main paper, except all gene sets compared have 500 genes instead of 100.

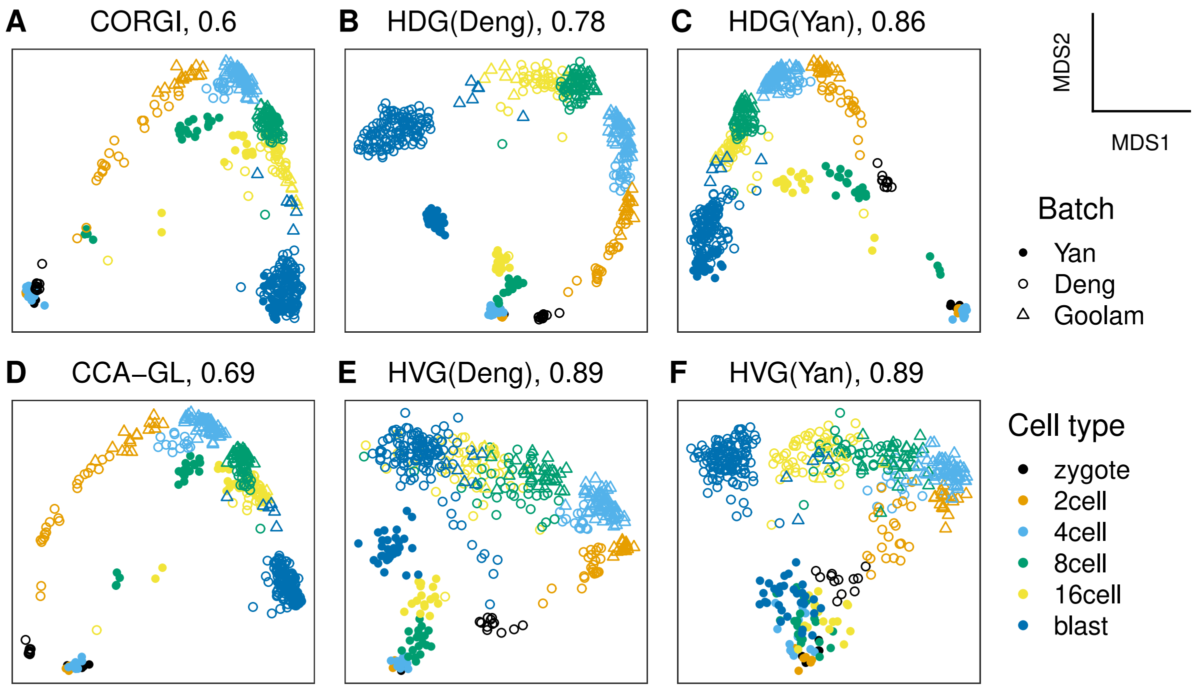


Figure S5: MDS on the embryogenesis datasets across various gene sets. Similar to Figure S4, except panel E and F show the cell embeddings using top 500 highly variable genes (HVG) instead of highly dropped-out genes. Each gene set contains 500 genes. Using highly variable genes (panel E and F) instead of highly dropped-out genes (panel B and C) resulted in lower quality.

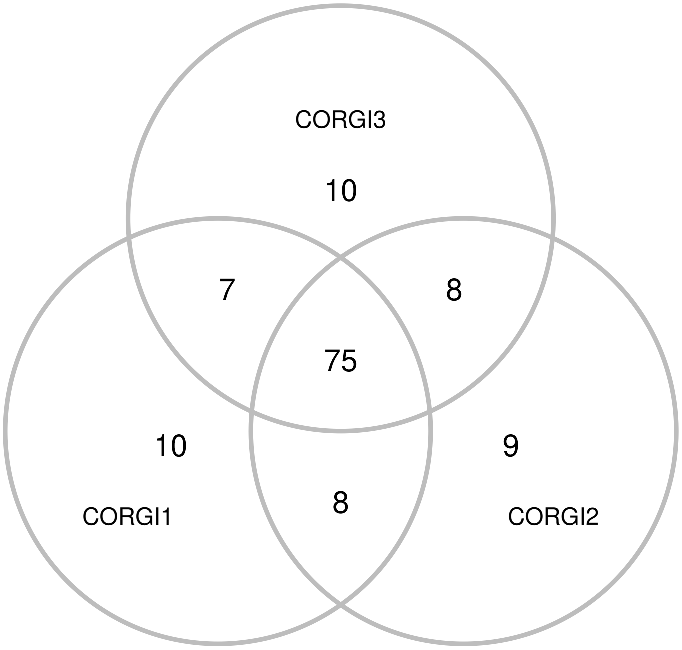


Figure S6: Venn diagram showing the top 100 genes from 3 independent runs of CORGI feature selection on the embryogenesis data.

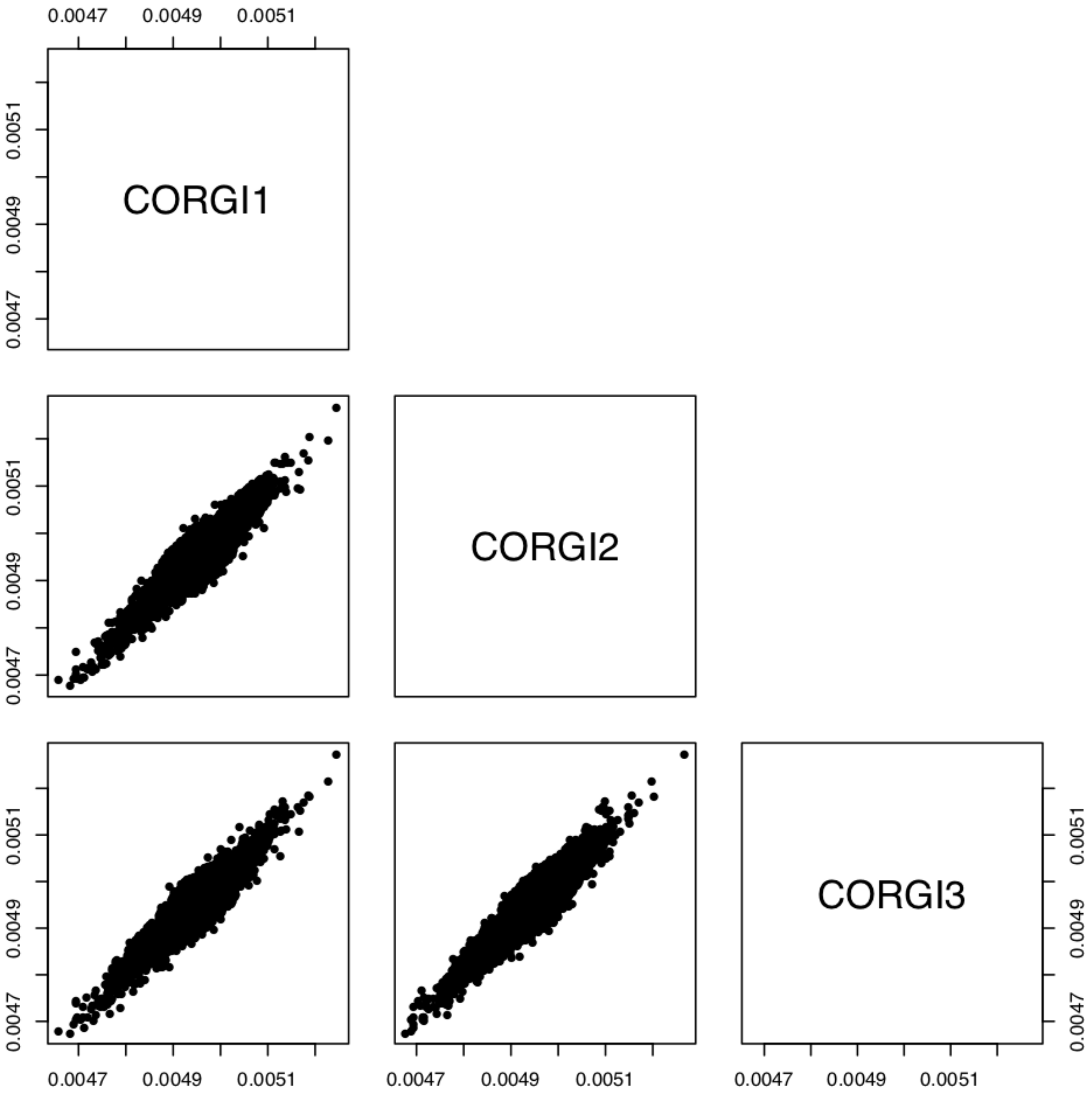


Figure S7: Pairwise plots of the principal curve scores between 3 independent runs of CORGI. Generated using the `pairs` function in the R base plot. Related to Figure S6.

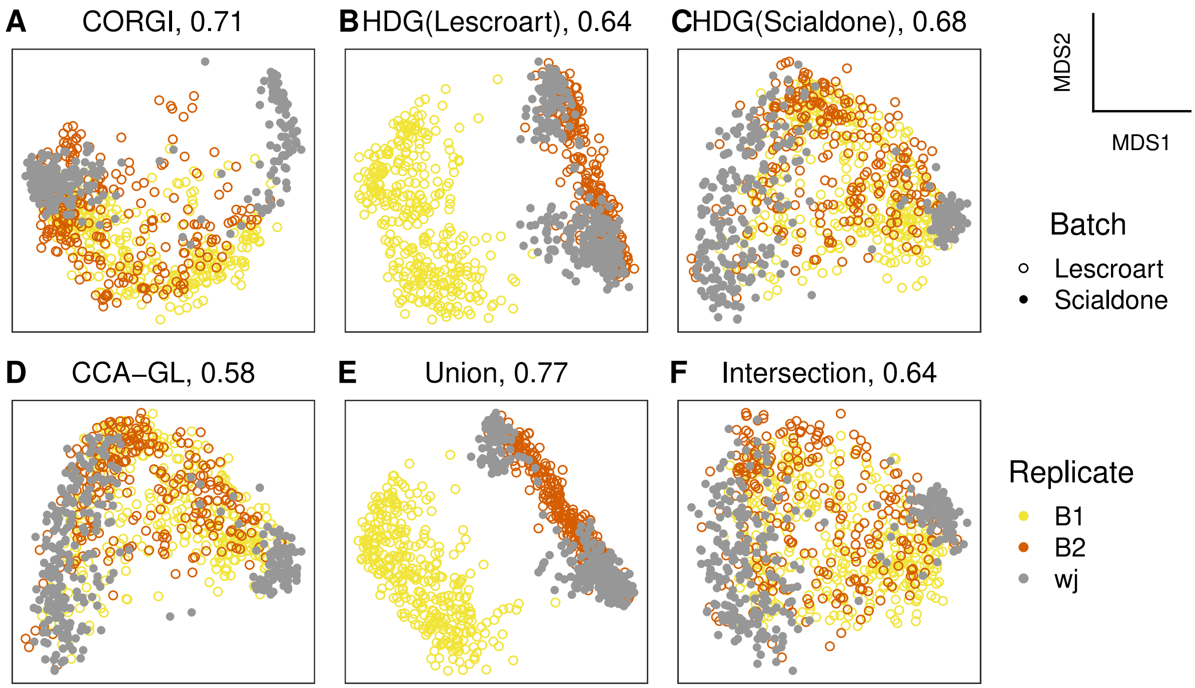


Figure S8: The cardiogenesis datasets colored by replicates. In the Lescroart batch, there is only one replicate while in the Scialdone batch, there are two.

### 3 Generating gene sets of a given size

Let $X_1$ and $X_2$ be the two batches of data. The method $HDG$ gives a ranking of genes and taking the top $n$ genes yields a gene set $HDG(X_i, n)$ of size $n$. In the main text, we suppress this parameter $n$ for brevity.

#### 3.1 Unions and intersections of highly dropped-out genes

Suppose now that we wish to pick $m$ such that $HDG(X_1, m) \cup HDG(X_2, m)$ has exactly $n$ genes. To find such an $m$, we define

$$f_1(m) = HDG(X_1, m) \cup HDG(X_2, m)$$

and use binary search to find $m$ such that $|f_1(m)| = n$. Such an $m$ may not necessarily exist since it is possible that our desired number of genes $n$ is skipped:

$$|f_1(m)| = n - 1, \qquad |f_1(m + 1)| = n + 1$$

Hence, we also consider

$$f_2(m) = HDG(X_1, m) \cup HDG(X_2, m + 1)$$

Clearly, there exists an $m$ such that $|f_1(m)| = n$ or $|f_2(m)| = n$. So for the union of HDGs, we set it as $f_1(m)$, or $f_2(m)$, whichever achieves the desired target size. The same method is used when the union is replaced by intersection.

#### 3.2 Augmenting a gene set by marker genes

Suppose we have a set of marker genes $M$. Suppose that $g$ is a function such that $g(m)$ is a gene set and $|g(m)| \leq |g(m + 1)|$ for all $m$. For example, $g(m) = f_1(m)$ as in the previous section. We wish to find $m$ such that $g(m) \cup M$ has a desired size, say $n$.

Consider

$$f(m) = g(m) \cup M.$$

Similar to the previous section, we apply binary search to find $m$ such that $|f(m)| = n$. Similar to the previous section, this isn't always possible, in which case, we find $m$ such that

$$|f_1(m) \cup M| = n \quad \text{or} \quad |f_2(m) \cup M| = n$$

where $f_1, f_2$ is defined as in the previous section.

### 4 List of gene sets

Below, genes in red are marker genes.

#### 4.1 Preimplantation embryogenesis

**CORGI**

SLC6A8 CD63 YARS TMPRSS2 ZSCAN5B ALDH18A1 WDR77 BTG4 ACCSL SPARC WDR69 NPM2 PRSS8 MAST3 GDAP1 SLC35E4 RSPO2 PADI6 ALPPL2 PLEKHG1 RASSF5 RELB ELF3 LRP2 ISG20L2 TOR1B PLBD1 UPP1 AURKC CCNO CXADR BMP15 TUBAL3 FHOD3 RPH3A SDC4 ANXA6 PGM1 DPPA4 GATA6 TDRD1 PABPC1L GATA2 FOLR4 UBTFL1 ZAR1L PSAT1 CRIP1 GPX4 GNA14 COMMD3 LHX8 BCL2L10 FAM167A PTGES PPP1R3D ALPL ADAMTSL1 ZP3 IGF2BP1 ZP2 LPCAT3 PFKFB3 TPP1 PARP12 FARP1 ENPEP MARCKSL1 SLC34A2 FBXL7 STMN3 FBXO31 SLCO3A1 AFAP1L2 ZFP42 MRPL12 BICC1 ASTL ENPP2 KLF6 AMPD3 HOXA7 TTC37 AQP3 MGST3 GTF2A1L IMPDH1 ZBTB16 RAB39 GCSH CDCA7L FBXO43 GEMIN4 KDM5B CCNB3 WEE2 AK7 TCEB2 CAPG RUNDC3B

**HDG(Deng)**

FBP2 TAGLN2 CITED1 CD63 ID2 KRT8 ALPPL2 GJB3 TFRC SDC4 CCNE1 PEMT ACAA2 SPIC NIN BHMT FABP3 GM2A LRP2 KRT18 VNN1 GLRX UPP1 ELF3 CSRNP2 CALCOCO2 TNFSF13B GSTA4 NT5C3L TSPAN8 SPP1 PSAT1 SLC46A1 PTGR1 CLDN4 GJB5 UQCRC1 ANXA2 HEXA LGALS1 MDH1 ID3 NPL PLIN2 KLF17 TDGF1 HSD17B14 ECH1 MGST3 PPT2 STMN3 LRRFIP1 HADH ENPEP GJA1 YIF1A FOLR1 LCP1 ABHD6 EMP2 CRY1 CLP1 CTSD SLC2A3 RIMKLB ASNS SOCS3 LRRC42 NUDT5 CTSZ DAP FBXL20 CCT6A XPNPEP1 PARP8 PCGF1 GSTO1 TMSB4X GSS ARHGEF16 SPARC NEDD4 DEF8 ACADVL CNDP2 SLC38A4 CLDN6 ZSCAN5B EPAS1 N6AMT2 ACSL4 ARHGAP29 SLC12A2 NAALAD2 ZFP57 CAPNS1 FAM46C CYB5R3 COMMD3 APOC1

**HDG(Yan)**

PLAC1L H1FOO LDHB GDF9 WEE2 OTX2 RGS2 PCP4L1 KPNA7 SLC2A3 ZP3 UCHL1 FAM151A PGAM1 DNMT1 PRAMEF6 S100A11 H2AFZ DNMT3L MYC ZP2 CCDC147 MFSD2A MBD3L2 WFDC2 PDE8B BPGM NPM2 GATA6 HIST1H2BK ZSCAN5B SLC6A5 S100A16 ALPPL2 SERPINF1 PLIN2 CTRB1 AURKC WBP5 ACCSL BIK OOEP ZAR1L HIST1H4C CNNM2 MSRA POLR3K KLF17 BCL2L10 FAM46C CXADR SOX15 BTG4 TUBB4 CD63 MBIP DTYMK IL17B PLEK2 ZIM3 ANXA2 DAB2 NEFM MAEL S100A10 ANKRD37 HABP2 ZFP42 S100A14 EID3 PADI6 HYLS1 SCO1 RPF2 GPR143 BASP1 PHOSPHO1 CSTB MRPL12 MRPL16 DUSP18 RARRES2 DEPDC7 MRPS17 MVP FAM13A CABYR RFK PATL2 SERTAD1 RBP7 CCNA1 HIST1H1A FBXO5 NDUFA8 TRIML2 CSRP2 ISG20L2 PXK PABPN1L

**CCA-GL**

BTG4 BCL2L10 ACCSL NDUFS6 RPL4 KRT18 ATP5E KRT8 USP2 JAG1 WDR69 GNA14 PAPD7 APOC1 NPM2

HSPE1 ACTB TOMM6 CPEB1 GSTP1 NLRP5 RSPO2 ATP5O CCT6A WEE2 DCLK2 COX4I1 ZBTB16 CCDC72 FYN ATP5G3 CCNO GNB2L1 TUBG2 CCNE1 PAIP1 PSMB1 DAAM1 RPH3A GDAP1 NDUFA4 BMP15 RPL36AL HSPD1 AKIRIN2 RNF38 PADI6 TUBAL3 ATP5F1 ATP5H ANXA2 TAF3 NHP2 TDRD1 PCGF1 DUSP7 MYL6 S100A10 JAZF1 ANXA7 CNOT6L TIAM1 SOX15 PPP1R3D ZSCAN5B EIF4A3 PSMA6 UQCR11 GATA3 IPO8 RPL22 UCHL1 GNG12 DCAF5 YPEL5 TAX1BP3 TULP3 TAGLN2 ZFAND5 UQCRH USP11 TMSB10 GPX4 PSMB3 EPC1 CLOCK CCNB1 BPGM RGS2 NDUFA8 COX8A ALKBH5 PSMB7 C1QBP TIMM13 SYCP2 TPM4 COX7A2 RPL32 PRDX1

**Union**

PLAC1L H1FOO LDHB GDF9 WEE2 OTX2 RGS2 PCP4L1 KPNA7 SLC2A3 ZP3 UCHL1 FAM151A PGAM1 DNMT1 PRAMEF6 S100A11 H2AFZ DNMT3L MYC ZP2 CCDC147 MFSD2A MBD3L2 WFDC2 PDE8B BPGM NPM2 GATA6 HIST1H2BK ZSCAN5B SLC6A5 S100A16 ALPPL2 SERPINF1 PLIN2 CTRB1 AURKC WBP5 ACCSL BIK OOEP ZAR1L HIST1H4C CNNM2 MSRA POLR3K KLF17 BCL2L10 FAM46C CXADR SOX15 FBP2 TAGLN2 CITED1 CD63 ID2 KRT8 GJB3 TFRC SDC4 CCNE1 PEMT ACAA2 SPIC NIN BHMT FABP3 GM2A LRP2 KRT18 VNN1 GLRX UPP1 ELF3 CSRNP2 CALCOCO2 TNFSF13B GSTA4 NT5C3L TSPAN8 SPP1 PSAT1 SLC46A1 PTGR1 CLDN4 GJB5 UQCRC1 ANXA2 HEXA LGALS1 MDH1 ID3 NPL TDGF1 HSD17B14 ECH1 MGST3 PPT2 STMN3

**Intersection**

SLC2A3 ZP3 UCHL1 WFDC2 NPM2 ZSCAN5B ALPPL2 PLIN2 OOEP KLF17 FAM46C CD63 ANXA2 S100A10 PADI6 CSTB SERTAD1 BMP15 RHEBL1 SLC34A2 LGALS1 CLIC1 MTHFD2 DDX3Y KLF11 PTPLAD1 PRSS8 YRDC KRT8 SLC6A8 ALG5 AIP PLA2G16 WDR47 WDR77 ALPL FAM83D APOC1 LTA4H PLBD1 PTGR1 CLDN4 PCGF1 SLC35E4 CRIP1 DAZL MRPL28 PTGES HHEX HTATIP2 FN1 GLRX EPAS1 GNG10 GOLM1 TIPARP ELF3 ZSWIM3 SERPINE2 YARS LMO7 IDH2 CLDN6 PARP12 UBTFL1 DUSP10 PLS3 FKBP6 STK31 SDC4 LPCAT3 SMPDL3A FABP3 PSAT1 KLHL21 CCT6A SNAI1 ZFYVE21 SLC1A3 AQP3 PFKFB3 RPL10L RAB20 NID1 SLC39A6 GSTO1 RALB AMOTL2 CAPRIN2 SLC16A6 GDF3 PEMT SPARC RNF130 AP4B1 TGFBRAP1 TRIB1 SLC25A13 SNUPN DET1

#### 4.2 Neurogenesis

Note: the 34 consensus genes analyzed in[2] are highlighted in red.

**CORGI**

Atp1a2 Apoe Ntsr2 Tnfrsf19 Slc1a3 Ppap2b Mfge8 Ndrg2 Hepacam Notch2 Dlx1 Ntrk2 Dtna Mmd2 Tspan7 Egfr Sp9 Tubb3 Cpe Dlx5 Mt2 Cst3 Sparcl1 Prnp Gm2a Itm2c Tiam2 Slc6a1 Qk Ednrb Ptn Ascl1 Ttyh1 S1pr1 D4Wsu53e Lmo3 Cd9 Sp8 Slc27a1 Cd81 Pdpn Sdc4 Ptprz1 Abracl Gas1 Sat1 Cspg5 Fgfr3 Ctsb Ckb Aldoc Soat1 Mt3 Miat Sox9 Basp1 Zfp36l1 Fam3c Sox8 F3 Tcf7l2 Notch1 Pitpnc1 Arhgap11a Atp1b2 Stil Dbi Btg2 Sorbs1 Tspan3 Ttk Gstm1 Gdap1 Cldn10 Arx Fbxo5

Lin7a Gstm5 Scrg1 Prdx6 Pdia3 Ska1 Mastl Myo1b Gpm6b Ube2ql1 Serpine2 Tmbim6 Itm2b Zfp78 Nav1 Spry2 Flrt2 Id2 Rcn2 Epha3 Acsl3 Elovl5 Laptm4a Slc25a13 Clu Ccnd2 Gja1 Tmsb4x Ptma Nono Htra1 Nrxn3 Dlx2 Jag1 Gm13826 Eef1a1 Igfbpl1 Sox11 Dlx6as1 Slc1a2 Rpl32 Malat1 Skil Bcan Set Marcks Tpx2 Plp1 Pdcd4 Actb Dcx Dpysl2 Glul Eif4a1 Rps11 Fabp7 Rps26 Mt1

**HDG(Dulken)**

Tubb5 Usp22 Egr1 Sept2 Zfp367 Ptp4a2 H3f3b Impact Fam120a Sdc4 Sparcl1 Lbr Golt1b Sept11 Top2a Twf1 Nras Lgr4 Ttyh1 Aldoc AI597468 Canx Tmpo Tmem55a Spin1 Cbx5 Efcab14 Ednrb Ppap2b Tial1 Sptbn1 Ntsr2 Slc1a3 Nfib Rrm2 Smc2 Klf13 Atp1a2 Klhl9 Dclk1 Id2 Rqcd1 Rrn3 Hn1 Tug1 Atrx Sat1 Nufip2 Uhrf1 Mmd2 Grm3 Paics Hsp90ab1 Poldip3 Cdk4 Map3k1 Gmps Clk1 Gsk3b Mfap2 Ssr1 Actr2 Gstm1 Fos Zfp266 2810474O19Rik Cd81 Ssr3 Prnp Cnbp Csde1 Cspg5 5430417L22Rik Hat1 Sh3bgrl Zscan26 Rdh13 Rpl4 Arl8b Rbm5 Smarce1 Map1b Mdh1 Spcs3 Ube2e3 Srsf7 Ilf3 Zfp655 Calm2 Abi2 Hnrnpa0 Cenpf Idh1 Inpp4a Ptges3 Sorbs1 Ccdc6 Arhgap5 Tpd52l2 Srpk2 Clu Ccnd2 Gja1 Tmsb4x Ptma Nono Htra1 Nrxn3 Dlx2 Jag1 Gm13826 Eef1a1 Igfbpl1 Sox11 Dlx6as1 Slc1a2 Rpl32 Malat1 Skil Bcan Set Marcks Tpx2 Plp1 Pdcd4 Actb Dcx Dpysl2 Glul Eif4a1 Rps11 Fabp7 Rps26 Mt1

**HDG(Llorens)**

Mcm7 Cdca7 Hn1 Ddx1 Tnfrsf19 Hdac2 Gnb2l1 Mta2 Nop56 Hes6 Cdk4 Dnajc9 Ppa1 Atf4 Sez6 2700094K13Rik Rrm2 Cops4 Stmn1 Sar1a Txnrd1 Coro1c Sumo3 Nasp Hnrnpr Rfc2 Tcp1 Hmgb3 Strap Fam210b Marcksl1 Ptbp1 Cdc16 Prps1 Smc3 Rpa2 Phip Vim Anp32b Cct2 Nap1l4 Kcne1l Gnai3 Cdc42se2 Sae1 Gars Actl6a Taf9 Tubg1 Odc1 Tnpo3 Psmc1 Ilf2 Tpm4 Dbn1 Hat1 Rsl1d1 Trim37 Cyb5b Eif4h Hnrnpab Fxyd6 Rbmx Mcm5 Mcm6 Dtymk Ywhah Kars Eif4a3 Rrm1 Polr2m Sf3b3 Klhdc3 Caprin1 Ncapd2 Med22 Pcna 2810417H13Rik Fxyd1 9530068E07Rik Ctnnb1 Arpc2 Nsg1 Papss1 Stt3a Abce1 Tubb5 Eprs Traf7 Drg1 Hibadh Cdc123 Kcnip3 G3bp1 Raf1 Slbp Smarca4 Acaa2 Anxa5 Mzt1 Clu Ccnd2 Gja1 Tmsb4x Ptma Nono Htra1 Nrxn3 Dlx2 Jag1 Gm13826 Eef1a1 Igfbpl1 Sox11 Dlx6as1 Slc1a2 Rpl32 Malat1 Skil Bcan Set Marcks Tpx2 Plp1 Pdcd4 Actb Dcx Dpysl2 Glul Eif4a1 Rps11 Fabp7 Rps26 Mt1

**CCA-GL**

Top2a Ccnd2 Tubb5 Ube2c Hmgb2 Tmsb4x Prc1 Gja1 Sox11 Rrm2 Clu Cdca2 Cst3 Ttyh1 Ckap2l Cdk1 Mt1 Ccnb1 Pbk Mt3 Mki67 Ptma Stmn1 2810417H13Rik Birc5 Ppap2b Tpx2 Cenpf Cdca8 Apoe Nusap1 Cenpa Spc25 Htra1 Atp1a2 Aldoc Ccna2 Fam64a Mis18bp1 H3f3b Slc1a2 Hn1 H2afz Cdca7 Sparcl1 Plk1 Kif22 Smc2 Tacc3 Pcna Uhrf1 2700094K13Rik Grm3 Racgap1 Hist1h4d Kif15 Dut Cenpe Dlx6as1 Kif11 Cdca3 Egr1 Casc5 Smc4 Hnrnpab Tubb3 Kif23 Esco2 Hmmr Tuba1b Slc1a3 Nasp Ckap2 Hat1 Dlx2 Ran Calm2 Rps11 Cspg5 Lig1 Npm1 Neil3 Aurka Rrm1 Cdk4 Cldn10 Dlx5

Malat1 Bub1b Bcan S1pr1 Glul Spc24 Ect2 Psrc1 Sp9 Rplp0 H2afx Rpl4 Ncapg Ranbp1 Rps9 Atp1b2 Marcksl1 Gstm1 Ntsr2 Ncapd2 Rpl8 Cdc20 Cd24a Hsp90ab1 Nop58 Nuf2 Arhgap11a Dcx Gpr37l1 Hmgb3 Nono Nrxn3 Jag1 Gm13826 Eef1a1 Igfbpl1 Rpl32 Skil Set Marcks Plp1 Pdcd4 Actb Dpysl2 Eif4a1 Fabp7 Rps26

**Union**

Tubb5 Usp22 Egr1 Sept2 Zfp367 Ptp4a2 H3f3b Impact Fam120a Sdc4 Sparcl1 Lbr Golt1b Sept11 Top2a Twf1 Nras Lgr4 Ttyh1 Aldoc AI597468 Canx Tmpo Tmem55a Spin1 Cbx5 Efcab14 Ednrb Ppap2b Tial1 Sptbn1 Ntsr2 Slc1a3 Nfib Rrm2 Smc2 Klf13 Atp1a2 Klhl9 Dclk1 Id2 Rqcd1 Rrn3 Hn1 Tug1 Atrx Sat1 Nufip2 Uhrf1 Mmd2 Grm3 Mcm7 Cdca7 Ddx1 Tnfrsf19 Hdac2 Gnb2l1 Mta2 Nop56 Hes6 Cdk4 Dnajc9 Ppa1 Atf4 Sez6 2700094K13Rik Cops4 Stmn1 Sar1a Txnrd1 Coro1c Sumo3 Nasp Hnrnpr Rfc2 Tcp1 Hmgb3 Strap Fam210b Marcksl1 Ptbp1 Cdc16 Prps1 Smc3 Rpa2 Phip Vim Anp32b Cct2 Nap1l4 Kcne1l Gnai3 Cdc42se2 Sae1 Gars Actl6a Taf9 Tubg1 Odc1 Tnpo3 Clu Ccnd2 Gja1 Tmsb4x Ptma Nono Htra1 Nrxn3 Dlx2 Jag1 Gm13826 Eef1a1 Igfbpl1 Sox11 Dlx6as1 Slc1a2 Rpl32 Malat1 Skil Bcan Set Marcks Tpx2 Plp1 Pdcd4 Actb Dcx Dpysl2 Glul Eif4a1 Rps11 Fabp7 Rps26 Mt1

**Intersection**

Tubb5 Egr1 Sdc4 Ttyh1 Tmpo Spin1 Rrm2 Rqcd1 Hn1 Atrx Uhrf1 Paics Poldip3 Cdk4 Mfap2 Hat1 Map1b Ilf3 Matr3 Mki67 Pbk Cpsf6 Cldn10 Dtymk Btg2 Timp3 Rrm1 Vbp1 Elavl4 Rbmx Hnrnpab Pak3 Sncaip Mcm4 Sumo3 Il18 Fubp1 Tceal8 Ivns1abp Tipin Rsl1d1 Eprs Cfl2 Bub3 Emc1 Nme1 Ptbp2 Trp53 Ascl1 Pdzd11 Lxn Zeb1 Xpo1 Smc3 Ankrd6 Tnfrsf19 Smc4 Ctnnb1 Tmem47 Rgcc Cct2 Tcf12 S100a1 Fkbp4 Fopnl Papola Prpf40a Maged1 Cdca8 Wrb Psmd14 Calu Thoc7 Eif4e Srsf6 Cnot7 Mrpl28 Nop56 Drg1 2810417H13Rik Actl6a Pbrm1 Cdc42se2 Smim7 Cep57 Dnaja2 Mgst1 Cct7 Zfp637 Slbp Gnb2l1 Tmem167 Rtn1 Tcp1 Nxf1 Ezh2 Miat Dpysl3 Psmd6 Sdhd Clu Ccnd2 Gja1 Tmsb4x Ptma Nono Htra1 Nrxn3 Dlx2 Jag1 Gm13826 Eef1a1 Igfbpl1 Sox11 Dlx6as1 Slc1a2 Rpl32 Malat1 Skil Bcan Set Marcks Tpx2 Plp1 Pdcd4 Actb Dcx Dpysl2 Glul Eif4a1 Rps11 Fabp7 Rps26 Mt1

#### 4.3 Cardiogenesis

##### CORGI

Cldn6 Pim2 Cdh1 Cldn7 Nodal Fgf3 Lhfp Kdr Esrp1 Fgf5 Chmp4c Slc7a3 Hand1 Fgf8 P3h2 Prdm6 Rab25 Utf1 Mt1 Bst2 L1td1 Asb4 Grb7 Nanog Wnt5a Eras Apln Pdpn Tor3a Tmem59l Pmp22 Phlda2 Psors1c2 Cxcl12 Mt2 Aplnr Rasgrp3 Trh Wnt2 Pdgfra Ccnb1ip1 Spint1 Ifitm1 Gldc Epha2 Nrp1

Pcsk5 Klhl6 Phldb2 Tpd52 Gstt2 Rbm47 Kcnk1 Capn2 Mesp1 Hoxb1 Tcea3 Nefl Tmem54 Zfp516 Cxcr4 Lama5 Jun Cap2 Heg1 1700013H16Rik 1700019D03Rik Wfdc2 Gfra3 Slc5a5 Galnt3 Epha4 Tcaf1 Vcan Grik3 Hs3st3b1 Myl7 Gata4 Slc35f2 Zfp36l3 Ppic Rspo3 Dppa5a Gsc H19 Greb1l Bspry Zfpm1 Glrx Sept8 Slc7a7 Pitx2 Crabp1 Hsf2bp Fads2 Foxf1 Epcam Psmb8 Inka1 Prickle1

**HDG(Scialdone)**

Tdgf1 Aplnr Xist Asb4 Kdr Msx2 Epcam Hand1 Snai1 Pim2 Ccnd2 Id1 Pdgfra Peg3 Fgf3 Gata4 Igfbp2 Nrp1 Fst Msx1 Pmp22 Ifitm1 Pcdh18 Cdh11 Otx2 Eomes Fgf5 Id2 Foxf1 Hoxb1 Dll3 Frzb Wls Bambi Cdh2 Lhfp Sgk1 Rspo3 Phldb2 Slc38a4 Mum1l1 Spint2 Myl7 Rasgrp3 Spin2c Wnt5a Pmaip1 Mixl1 Etv5 Nkd1 Laptm4b Dusp6 Prdm6 Crabp1 Wrb Cfc1 Wfdc2 Atp6v1b2 Ucp2 Cldn6 Mcm5 Ccn1 Jun Prps2 Epha4 Mid1ip1 Mt2 Dkk1 Pcdh19 Stx3 Kcnk1 Ubr7 T Glipr2 Bmp4 Crip2 Gnb4 Capn2 Greb1l Nfil3 Zic3 F11r Bcat2 Cacfd1 Sp5 Tmem185a Crmp1 Colec12 Epb41l3 Rragd Fos Cd276 Ina Cnn2 Tbx3 Igf2 Ap1m2 Xab2 Dll1 Gna14

**HDG(Lescroart)**

Foxf1 Hoxd13 Sox3 Vax1 Bicd2 Irx3 Six3 Tdgf1 Fgf3 Pcdh19 St3gal2 Pou3f3 Mixl1 Mest Ncoa1 Kcnk12 Dusp15 Slc4a4 Lhfp Epb41l3 Xist Foxe1 Mesp1 Glipr2 Phldb2 Kdr Dll1 Kcnc4 Fgf8 Dll3 Lefty2 Eomes Klhl6 Pim2 Tenm3 Cyp26a1 Evx1os Aplnr Nin Capn2 St6galnac4 Nnat Asb4 Msx1 Plbd2 Rasgrp3 Pdgfra Epha4 Tdgf1-ps1 Slc7a7 Gsc Wnt3 Sept8 Bag2 Crmp1 Zxda Bmp7 Evx1 Camk2n1 Ap1m2 Fgf15 Mycl Lzts1 Crabp1 Epha1 Ankrd54 Ndn Slc52a2 Inka1 Anxa5 Ucp2 Def8 Dusp6 Nectin2 Mfge8 Elovl1 Cldn12 Gata6 Trp53i11 Chst7 Ascl1 Prkcd Psd3 Chtf18 Rps2-ps13 Mutyh Mapk7 Zfp553 Pdlim2 Dhx38 Cdc42ep4 Ext2 Soga3 Msx2 Tollip Tspan13 Hic1 Fhl1 Tmem185a 1700086L19Rik

**CCA-GL**

Ifitm1 Phlda2 Ube2c Epcam Pim2 Dnmt3b Hand1 Fgf8 Igf2 Myl7 Krt18 Bmp4 H19 Cldn6 Bex1 Pmp22 Laptm4b Pkm Foxc2 Crabp1 Sfrp1 Fn1 Dusp9 Igfbp4 Msx2 Pou5f1 Ppic Fscn1 Ccnb1 Krt8 Lef1 Plk1 Gm9843 Nanog Gpc3 Psme2b Spint2 Sub1 Cdh1 Stard8 Ccnd2 Ung Mest Cldn7 Prdm6 Lhfp Aurka Apoe Epb41l3 Ndufa4 Prtg Dok4 Tmem88 Dll1 Prc1 Fst Chchd10 Dll3 Kdr Morc4 Cox6b1 Hsd11b2 Hmga2 Top2a Sparc Nop10 Atpif1 Psme2 Rbms1 Rps5 Capn6 Ahnak Aplnr S100a10 Eif5a Hoxb1 Utf1 Tbx3 Tdgf1 Spin2c Cox7a2 Lefty2 Nrp1 Rps20 Mesp1 T Peg3 Marcksl1 Igfbp2 Foxh1 Grb7 Apln Slc7a3 Polg Nodal Cox8a Rbpms2 Cst3 Tbx6 Notch1

**Union**

Foxf1 Hoxd13 Sox3 Vax1 Bicd2 Irx3 Six3 Tdgf1 Fgf3 Pcdh19 St3gal2 Pou3f3 Mixl1 Mest Ncoa1 Kcnk12 Dusp15 Slc4a4 Lhfp Epb41l3 Xist Foxe1 Mesp1 Glipr2 Phldb2 Kdr Dll1 Kcnc4 Fgf8 Dll3 Lefty2 Eomes Klhl6 Pim2 Tenm3 Cyp26a1 Evx1os Aplnr Nin Capn2 St6galnac4 Nnat Asb4 Msx1

Plbd2 Rasgrp3 Pdgfra Epha4 Tdgf1-ps1 Slc7a7 Gsc Wnt3 Sept8 Bag2 Crmp1 Zxda Bmp7 Evx1 Msx2 Epcam Hand1 Snai1 Ccnd2 Id1 Peg3 Gata4 Igfbp2 Nrp1 Fst Pmp22 Ifitm1 Pcdh18 Cdh11 Otx2 Fgf5 Id2 Hoxb1 Frzb Wls Bambi Cdh2 Sgk1 Rspo3 Slc38a4 Mum1l1 Spint2 Myl7 Spin2c Wnt5a Pmaip1 Etv5 Nkd1 Laptm4b Dusp6 Prdm6 Crabp1 Wrb Cfc1 Wfdc2 Atp6v1b2

**Intersection**

Foxf1 Tdgf1 Fgf3 Pcdh19 Mixl1 Lhfp Epb41l3 Xist Glipr2 Phldb2 Kdr Dll1 Fgf8 Dll3 Lefty2 Eomes Klhl6 Pim2 Cyp26a1 Aplnr Capn2 St6galnac4 Asb4 Msx1 Rasgrp3 Pdgfra Epha4 Slc7a7 Wnt3 Sept8 Bag2 Crmp1 Bmp7 Evx1 Ap1m2 Fgf15 Mycl Crabp1 Epha1 Ndn Ucp2 Def8 Dusp6 Elovl1 Cldn12 Gata6 Prkcd Cdc42ep4 Msx2 Tspan13 Tmem185a Prps2 Flnb Fos Fbxo21 Dpysl5 Cfc1 Gfpt2 Myl7 Nfil3 Cenpl Dapk1 Coq9 Pmaip1 Frmd8 Crip2 Prc1 Gna14 Swap70 Wdr6 Jun Dkk1 AI597479 Dusp1 Hs3st3b1 1810055G02Rik Ctsc S1pr2 Tbx3 Bid Zfp110 Ccdc71 Abhd4 Tuba4a Wnt5a Gnb4 Wnt2 Ina Rassf7 Ezr Pgm1 Tenm4 Fgf5 Hmox1 Nkd1 Tmem184c Hdac6 Tnfrsf19 Mboat2 Prdm6
